# Efficient Estimation of Large-Scale Spatial Capture-Recapture Models

**DOI:** 10.1101/2020.05.07.081182

**Authors:** Daniel Turek, Cyril Milleret, Torbjørn Ergon, Henrik Brøseth, Perry de Valpine

## Abstract

Capture-recapture methods are a common tool in ecological statistics, which have been extended to spatial capture-recapture models for data accompanied by location information. However, standard formulations of these models can be unwieldy and computationally intractable for large spatial scales, many individuals, and/or activity center movement. We provide a cumulative series of methods that yield dramatic improvements in Markov chain Monte Carlo (MCMC) estimation for two examples. These include removing unnecessary computations, integrating out latent states, vectorizing declarations, and restricting calculations to the locality of individuals. Our approaches leverage the flexibility provided by the `nimble` R package. In our first example, we demonstrate an improvement in MCMC efficiency (the rate of generating effectively independent posterior samples) by a factor of 100. In our second example, we reduce the computing time required to generate 10,000 posterior samples from 4.5 hours down to five minutes, and realize an increase in MCMC efficiency by a factor of 25. We also explain how these approaches can be applied generally to other spatially-indexed hierarchical models. R code is provided for all examples, as well as an executable web-appendix.

## 1 Introduction

Capture-recapture methods are primary tools for estimating abundance and demographic parameters in populations. These methods model longitudinal encounter histories of individuals in a population. Spatial capture-recapture (SCR) models account for individual and trap-specific capture probabilities depending on individuals’ latent centers of activity and space-use in relation to the explicit location of traps or other detectors (Efford, 2004; Borchers and Efford, 2008). Closed SCR models provide more precise and robust estimates of population densities than non-spatial models, and also enable estimation of the spatial distribution of individuals and associated parameters.

Despite their popularity, SCR models encounter numerous computational challenges which pose serious obstacles for their practical use (Gardner et al., 2018). For large study areas with many detectors, determining the probability of a capture history becomes very computationally costly because it involves calculations for all detectors, which is problematic for large-scale studies (Milleret et al., 2018b). Modeling the movement of activity centers often induces inefficient MCMC updating, as do methods for imposing spatial constraints on activity center locations. And data augmentation of never-observed individuals can lead to unnecessary calculations.

Bayesian hierarchical models, such as SCR models, are often formulated using the BUGS modeling language (Lunn et al., 2009) and estimated using Markov chain Monte Carlo (MCMC; Brooks et al., 2011). Mainstream MCMC software includes WinBUGS, JAGS (Plummer, 2003), and Stan (Stan Development Team, 2014). Recently, the `nimble` R package has been developed, offering new degrees of customization for MCMC (de Valpine et al., 2017). Custom-written distributions and the flexibility of `nimble`’s MCMC system have provided substantial improvements in non-spatial capture-recapture models (Turek et al., 2016) and the study of MCMC algorithms (Turek et al., 2017).

We use `nimble` to demonstrate several generally applicable techniques for improving MCMC efficiency of (1) a simple but computationally-intense SCR model (Milleret et al., 2019), and (2) an open robust-design SCR model (Ergon and Gardner, 2014). We increase MCMC efficiency by vectorizing calculations, applying custom MCMC sampling strategies, implementing model-specific likelihood calculations, disabling unnecessary model calculations, and restricting trap calculations to the locality of each individual. Using these techniques, we achieve efficiency gains of a factor of 100 in the first example and a factor of 25 in the second example.

## 2 Materials and Methods

We consider two example SCR models which both present computational challenges. The first (“Wolverine”) considers a simple closed SCR model for data from non-invasive genetic sampling of wolverines on a large spatial scale in Norway (*Gulo gulo*, Milleret et al., 2019). The second (“Vole”) is a more complex SCR model on a smaller spatial scale, modeling an open population of field voles with activity-center movements (*Microtus agrestis*, Ergon and Gardner, 2014). We first describe each model, followed by the strategies used to improve MCMC efficiency. Finally, we describe the metric used to measure MCMC efficiency.

### 2.1 Wolverine Model

This example has a spatial extent over 200,000 km^2^. The data, collected throughout Norway, consist of 453 detections from 196 individually identified female wolverines using noninvasive genetic sampling and search encounter methods (Milleret et al., 2019). The search area was discretized to a detector grid with a 2km resolution, and only searched grid cells were included in the analysis. This resulted in 17,266 unique detectors, with binary-valued detections of individuals within grid cells. Data and additional details are available at the dryad repository (Milleret et al., 2018a).

The Wolverine model combines a spatial point process model of individual activity centers (ACs), data augmentation to model the true population size, and an observation model for detection probabilities and capture histories. Define the AC of individual *i* as 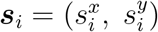, where 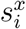 and 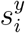 follow independent uniform prior distributions spanning the study area. As some regions are unsuitable habitat (*i.e.*, water), AC locations must be constrained. We use a habitat mask by defining a binary matrix ***H*** over the study area, where *H*_*x,y*_ = 1 indicates that cell (*x, y*) is suitable habitat. AC locations are then constrained as 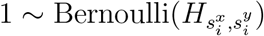, where 1 is a unit data value.

For data augmentation (Royle, 2009), we add *N*_aug_ virtual individuals. The augmented matrix *y* has dimension (*N*_obs_ + *N*_aug_) × *R*, with *R* = 17, 266 detectors and *N*_obs_ = 196 unique individuals. Define binary variables *z*_*i*_ with independent *z*_*i*_ ~ Bernoulli(*ϕ*) prior distributions, representing inclusion in the population. For the *N*_obs_ sighted individuals, *z*_*i*_ = 1 is observed data, while the remaining *z*_*i*_ are unobserved. Total population size *N* is estimated as 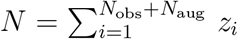, using the prior distribution *ϕ* ~ Uniform(0, 1) to induce a flat prior on *N* (Royle et al., 2007).

The probability of detecting individual *i* at detector *r* is 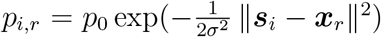, where ***x***_*r*_ is the location of detector *r* and *p*_0_ and *σ* are the maximal and scale of decay for detection probability. Detections are modeled as *y*_*i,r*_ ~ Bernoulli(*p*_*i,r*_ *z*_*i*_). The complete Wolverine model definition is given in (1), where indices *r* take the range 1, …, *R*.

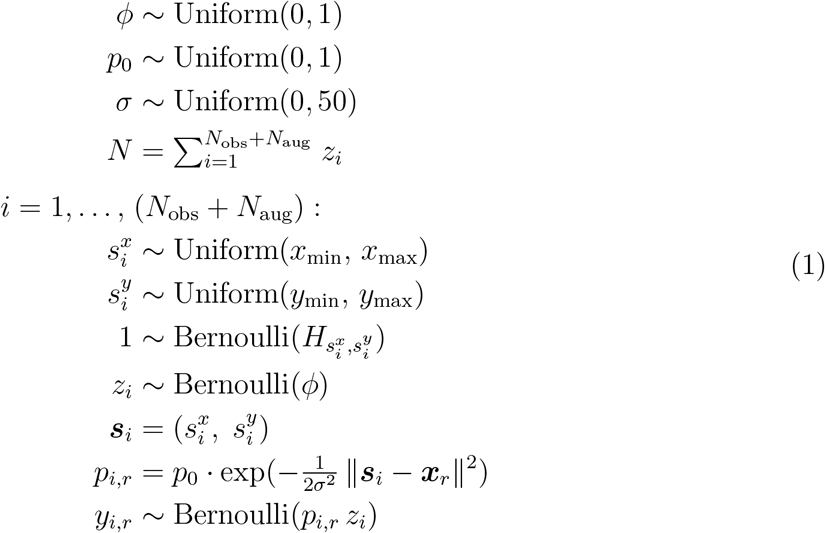

We use four refinements of the model and MCMC sampling, with the goal to improve MCMC efficiency: (1) Vectorize computations and put the habitat mask into a custom distribution, (2) jointly sample AC components, (3) restrict calculations to local detectors and sparse representation of data, and (4) skip unnecessary calculations when *z*_*i*_ = 0. We next describe each of these techniques, and `nimble` code corresponding to each cumulative refinement appears in Appendix A.

#### 2.1.1 Vectorized Computations

Vectorization refers to carrying out a set of matching model computations more efficiently, as is possible in `nimble` but neither WinBUGS or JAGS. `nimble` supports vectorized model declarations, reducing the total nodes in the model and potentially improving MCMC efficiency. We vectorized both detection probabilities and data likelihoods for each individual across the *R* detectors. For the vector of detection probabilities ***p***_*i*,1:*R*_, we used a vectorized model declaration. For the vectorized data likelihood of ***y***_*i*,1:*R*_, we used a custom likelihood function for the entire (length-*R*) observation history of one individual.

This technique is only beneficial when the *entire* joint likelihood of ***y***_*i*,1:*R*_ is always calculated simultaneously, as is the case here for updates of *p*_0_, *σ*, or *z*_*i*_. In a different model, this technique could result in inefficiencies if any MCMC updates require likelihood calculation for only a subset of ***y***_*i*,1:*R*_.

#### 2.1.2 Joint Sampling of AC Locations

We apply joint (block) sampling of the 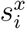 and 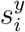 coordinates of each AC. `nimble` allows the assignment of block samplers to arbitrary variables, applying multi-dimensional Metropolis-Hastings sampling. This results in computational savings since an MCMC update of ***s***_*i*_ requires only one calculation of all (length-*R*) relevant detection probabilities and data likelihoods. In contrast, independent updates of the 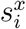 and 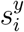 components will require two likelihood evaluations, one for each component.

#### 2.1.3 Local Detector Evaluations and Sparse Observation Matrix

We move detection probability calculations inside the vectorized likelihood, and additionally restrict these calculations to detectors within a maximum realistic radius (*d*_max_) of the AC ***s***_*i*_. In advance, we identify the set of detectors located within *d*_max_ from each cell of the habitat matrix. The modified distribution identifies the grid cell containing ***s***_*i*_, and the set of detectors within *d*_max_ from it. Calculations of *p*_*ir*_ are then restricted to this set of detectors.

We also convert to a sparse representation of the detection matrix *y*. In this representation, each row contains the detector identification numbers (values of *r*) that detected one individual. The number of columns is therefore equal to the maximum number of detections of any particular individual. This sparse representation allows for a smaller model and equivalent, but more efficient, likelihood calculations.

#### 2.1.4 Skip Unnecessary Calculations

Calculations can be avoided when any *z*_*i*_ = 0, that is, an augmented virtual individual is not currently included in the population. In that case, neither the distances to each detector nor the detection probabilities need be calculated. We modify the custom likelihood again, to accept *z*_*i*_ as an argument. When *z*_*i*_ = 1, the calculations take place as before. When *z*_*i*_ = 0, the likelihood is one if the individual was never observed – always the case for augmented individuals – which can be calculated without any distances or detection probabilities. This modification can save substantial computation, especially when *N*_aug_ is large, that being the conservative approach.

### 2.2 Vole Robust-Design Model

Our second example considers a robust-design SCR model of field voles in the Kielder Forest of northern England (*Microtus agrestis*, Ergon and Gardner, 2014), with four primary sampling occasions and nested secondary trapping sessions. A total of 158 unique individuals are considered to have static ACs within primary occasions, but to disperse between primary occasions. See Ergon and Gardner (2014, Appendix S2) for further details, (Ergon and Lambin, 2013) for the data, and Appendix B.1 for the original JAGS code.

The Vole model contains individual survival between primary sampling occasions, dispersal of ACs between primary occasions, and spatial capture-recapture from capture histories. Define the AC of individual *i* on primary occasion *k* as 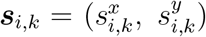. On first capture, the components 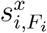 and 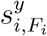 are given uniform prior distributions spanning the mean location of captures during that occasion. The dispersal between primary occasions *k* and *k* + 1 uses a uniformly-distributed dispersal angle *θ*_*ik*_, and an exponentially-distributed dispersal distance *d*_*ik*_ with rate parameter 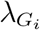, where *G*_*i*_ is the sex of individual *i* (1: female; 2: male), and λ_1_ and λ_2_ are sex-specific parameters. Thus, the AC components are related across primary occasions as 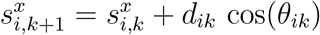 and 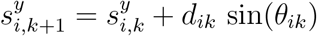.

The survival model uses binary indicator variables, where *z*_*i,k*_ = 1 indicates individual *i* is alive on occasion *k*. We condition on the first observation in primary occasion *F*_*i*_, as 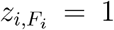. The survival process follows as 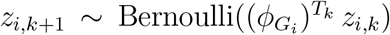, where survival probability depends on sex and temporal duration. *G*_*i*_ gives the sex of individual *i*, *T*_*k*_ is the time (in months) between occasions *k* and *k* + 1, and *ϕ*_1_ and *ϕ*_2_ are sex-specific survival rates. When 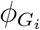 is a function of a continuous covariate, the model is only invariant to the choice of time unit of *T*_*k*_ when using a loglog (log-hazard) link (Ergon et al., 2018).

The observation model uses hazard rates to calculate trap capture probabilities. For individual *i*, on secondary trapping session *j* of primary occasion *k*, the capture hazard rate 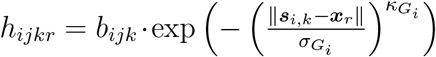, where the location of trap *r* is ***x***_*r*_, and each *κ*_*j*_ and *σ*_*j*_ are sex-specific observation parameters. Baseline hazard is 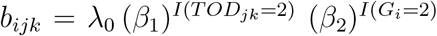, using indicator function *I*(·), time of day *TOD*_*jk*_ (1: evening; 2: morning), and baseline hazard rate λ_0_. *β*_1_ is the effect of morning trapping sessions, and *β*_2_ is that of males.

Total capture hazard rate is 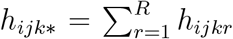. Probability of “no capture” is *π*_*ijk*0_ = exp (−*h*_*ijk*∗_ *z*_*i,k*_), which is unity when *z*_*i,k*_ = 0. Probability of capture is 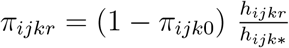 in trap *r*, accounting for competing risks among traps and satisfying 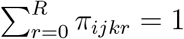.

The “ones trick” is used to induce the correct likelihood calculation. Observation data *y* is a 3-dimensional array, where *y*_*ijk*_ = 0 indicates that individual *i* was not captured in trapping session *j* of primary occasion *k*, and *y*_*ijk*_ = *r* indicates a capture in trap *r*. The complete Vole model definition is given in (2), where all indices *j* take the range of the number of secondary trapping sessions in the relevant primary occasion *k*, and all indices *r* assume the range 1, *…, R*.

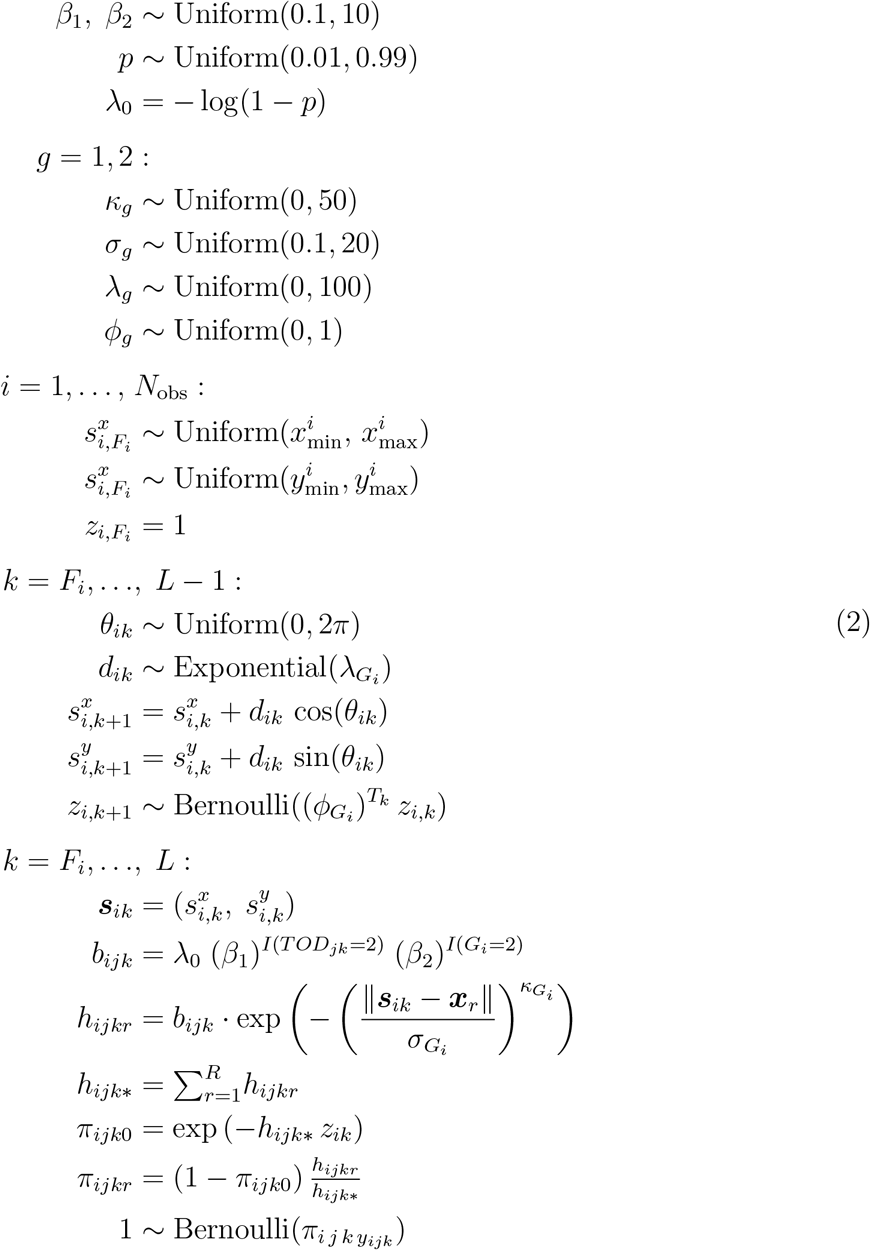

We apply three cumulative refinements to the model and MCMC sampling: (1) Jointly sample correlated dimensions and marginalize over *z*_*i*_ indicator variables, (2) use a custom bivariate dispersal distribution, and (3) restrict trap calculation to the vicinity of each AC. Next we describe these techniques, and nimble code corresponding to each appears in Appendix B.

#### 2.2.1 Joint Sampling and Marginalization

We apply joint samplers for updating two pairs of parameters: {*κ*_1_, *σ*_1_} and {*κ*_2_, *σ*_2_}, as these pairs each determine the trap hazard rates for one sex. Trial runs confirm that these pairs exhibit high posterior correlation, so we expect block samplers will improve mixing.

Next, we integrate (marginalize) over the latent *z*_*i,k*_ indicator variables to directly calculate the unconditional likelihood of capture histories. This reduces the model size and the dimension of sampling, and can improve MCMC mixing since parameter updates are no longer conditional on the “current” values of each *z*_*i,k*_. This is done in `nimble` using a custom likelihood. This calculation is a finite summation over the possible *z*_*i,k*_ states, similar to the filtering employed in Turek et al. (2016, Section 2.3.2). When individuals are known to be alive (up to the final capture), the likelihood is survival multiplied by the probability of the observed capture history. Subsequent to the final capture, forward-filtering is used to calculate the likelihood of the remaining non-capture events, accounting for uncertainty in survival.

#### 2.2.2 Custom Dispersal Distribution

We originally modeled dispersal distances and angles as random variables subject to MCMC sampling, a standard approach for movement models. This results in high computational cost because any proposed update to dispersal distance or angle (especially for *early* primary occasions) results in a large chain of calculations to determine the updated ACs, detection probabilities, and detection likelihoods *for all subsequent occasions*. Specifically, say we make an MCMC proposal for modifying *d*_11_, the dispersal distance for the first individual, between the first and second primary occasions. This MCMC update will require re-evaluating each ***s***_1,2_, ***s***_1,3_, …, ***s***_1,*L*_, up through the AC of the final primary occasion. Further, detection probabilities and data likelihoods for each AC also need be recalculated.

We reparameterize this model using a custom distribution of activity center locations that is induced by the distributions of turning angle and distance, as 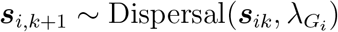. This distribution is centered around the current AC and is mathematically equivalent to the original {*d, θ*} parameterization. Now, updates of ***s***_*i,k*_ do *not* induce a large chain of ensuing calculations, but rather, only the likelihoods corresponding to ***s***_*i,k*_ and ***s***_*i,k*+1_ must be calculated. The custom distribution is given by 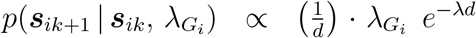, where *d* = ‖***s***_*ik*+1_ − ***s***_*ik*_‖, and omitting constants of proportionality which are not necessary for sampling. We recognize 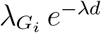 as the exponential density for the dispersal distance *d*. The factor of 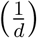 results from the Jacobian term in the change-of-variables between polar and Cartesian coordinates. From an implementation standpoint, these density calculations take place on a logarithmic scale. This technique can be similarly applied when using other distributions for dispersal distance, by substituting in the density of the alternate dispersal distribution.

#### 2.2.3 Local Trap Calculations

During MCMC sampling, the capture hazard rate *h*_*ijkr*_ and associated likelihood terms are calculated for all traps, regardless of an individual’s current AC. This is inefficient, since when ***s***_*i,k*_ is “far” from a trap *r*, then *h*_*ijkr*_ will be extremely low, and a capture in trap *r* is exceedingly unlikely. Its contribution to 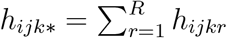 is negligible, as is the probability of capture in trap *r*. The original BUGS modeling language lacks the ability to conditionally disable calculations, and hence all the capture hazard rates must always be computed.

We introduce logic such that *h*_*ijkr*_ is only calculated for traps within a distance *d*_min_ of the individual’s AC. For traps located further, we assign *h*_*ijkr*_ a small positive value. This will not affect the sum *h*_*ijk∗*_, but still allows for a non-zero probability of capture. Here, we let *h*_*ijkr*_ = 10^−14^ for traps outside a radius *d*_min_ = 40 from each individual’s AC.

We introduce a discretized grid over the study area, and pre-compute indices of traps within a radius *d*_min_ from each grid cell. Using this, a custom `nimble` function returns the indices of “local traps” nearest to any ***s***_*i,k*_, and subsequently calculates hazard rates only for the “local” traps. This is similar to the local trap calculations used in the previous example, but implementations are different on account of the discretized habitat mask used there.

### 2.3 MCMC Efficiency

We define MCMC efficiency as the number of effectively independent posterior samples produced per second of MCMC runtime (excluding upfront time of model building and compilation). Distinct model parameters will typically mix at different rates, thus having distinct posterior effective sample sizes (ESS), and therefore a distinct measure of efficiency. In addition to presenting the MCMC runtimes and MCMC efficiency of all model parameters, we also summarize performance using the minimum and mean efficiencies among all model parameters. This definition of efficiency captures the tradeoff between quality of mixing and computational speed. Some algorithms may mix slowly (producing a low ESS) but execute sufficiently fast that they achieve high efficiency. Other algorithms may mix quickly (producing a high ESS) but require significantly longer execution time and thus achieve low efficiency.

## 3 Results

Here we describe the performance resulting from each formulation or sampling strategy of the Wolverine and Vole example models. All algorithm runtimes, ESS estimates, and MCMC efficiencies reflect independent chains of 10,000 posterior samples. We do not present the posterior inferences (*e.g.,* posterior mean, median, etc.), as they are qualitatively identical to the original published analyses.

### 3.1 Wolverine Model

We assess performance of the Wolverine model using total population size (*N*), probability of detection (*p*_0_), and scale factor (*σ*). Results for the four stages of iterative improvement described in Section 2.1 will be denoted as Nimble1 (vectorization), Nimble2 (joint sampling), Nimble3 (evaluating local detectors), and Nimble4 (skipping unnecessary calculations).

As in Milleret et al. (2019), the JAGS model was unable to complete, crashing after 30 days. Transitioning to `nimble` considerably reduced memory usage and runtime, as we fit the Nimble1 model in 26 hours. ESS values were in the range of 100 to 200 for all parameters, indicating high posterior auto-correlation. In combination with the long runtime, this produced MCMC efficiencies on the order of 10^−3^ for all parameters. The addition of joint sampling in the Nimble2 version decreased runtime to 20 hours. Parameter ESS values were similar to the Nimble1 model, giving a small improvement in efficiency.

We observed major improvement in the Nimble3 version, using the local trap evaluations and a sparse representation of the observation matrix. MCMC runtime reduced to 30 minutes, by a factor of 40 relative to the Nimble2 model. As we expect, ESS values were unchanged, and the resulting MCMC efficiencies were in the range of 0.1 to 0.3 (Figure 1).

**Figure 1:**
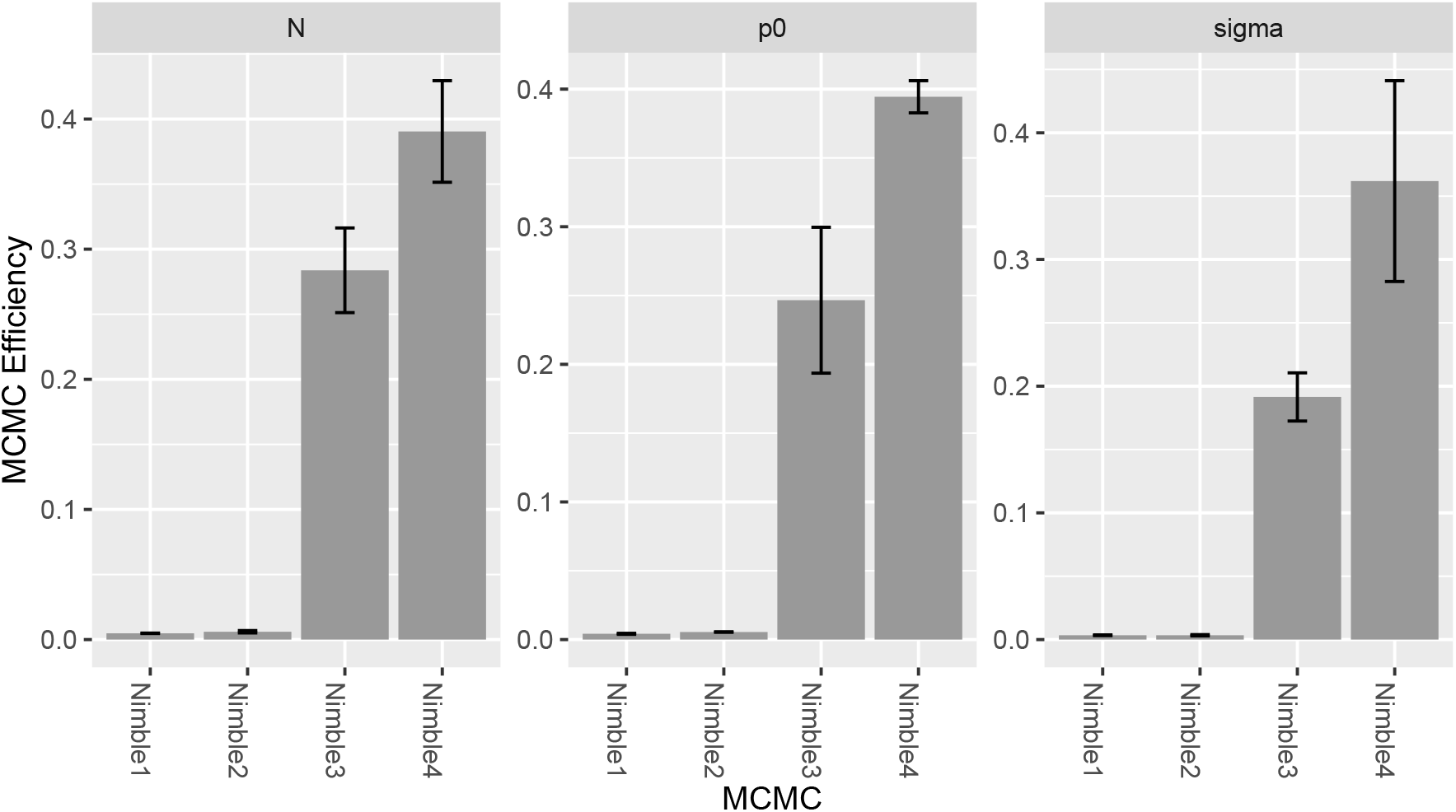
MCMC efficiency for the Wolverine example, for all parameters and model formulations. Efficiency values are averaged over three independent chains, error bars showing standard deviation.

The Nimble4 version, disabling unnecessary model calculations, reduced MCMC runtime by an additional factor of two, down to 16 minutes. Accordingly, MCMC efficiencies increased by nearly a factor of two. Relative to the initial Nimble1 formulation, we have achieved increases in both the minimum and mean efficiencies of 100-fold. Concretely, while it was not even possible to fit the original version of this model using JAGS, the initial Nimble1 formulation would require 3.5 days to generate 1,000 ESS for all parameters, and the final Nimble4 model can accomplish the same in 51 minutes.

Figure 2 presents the minimum and mean efficiencies across all model parameters for each formulation of the Wolverine model, and all results for the Wolverine example appear in Table 4. An executable version of the Nimble4 Wolverine model is available at the web-appendix http://danielturek.github.io/public/scr/wolverine_example.html.

**Figure 2:**
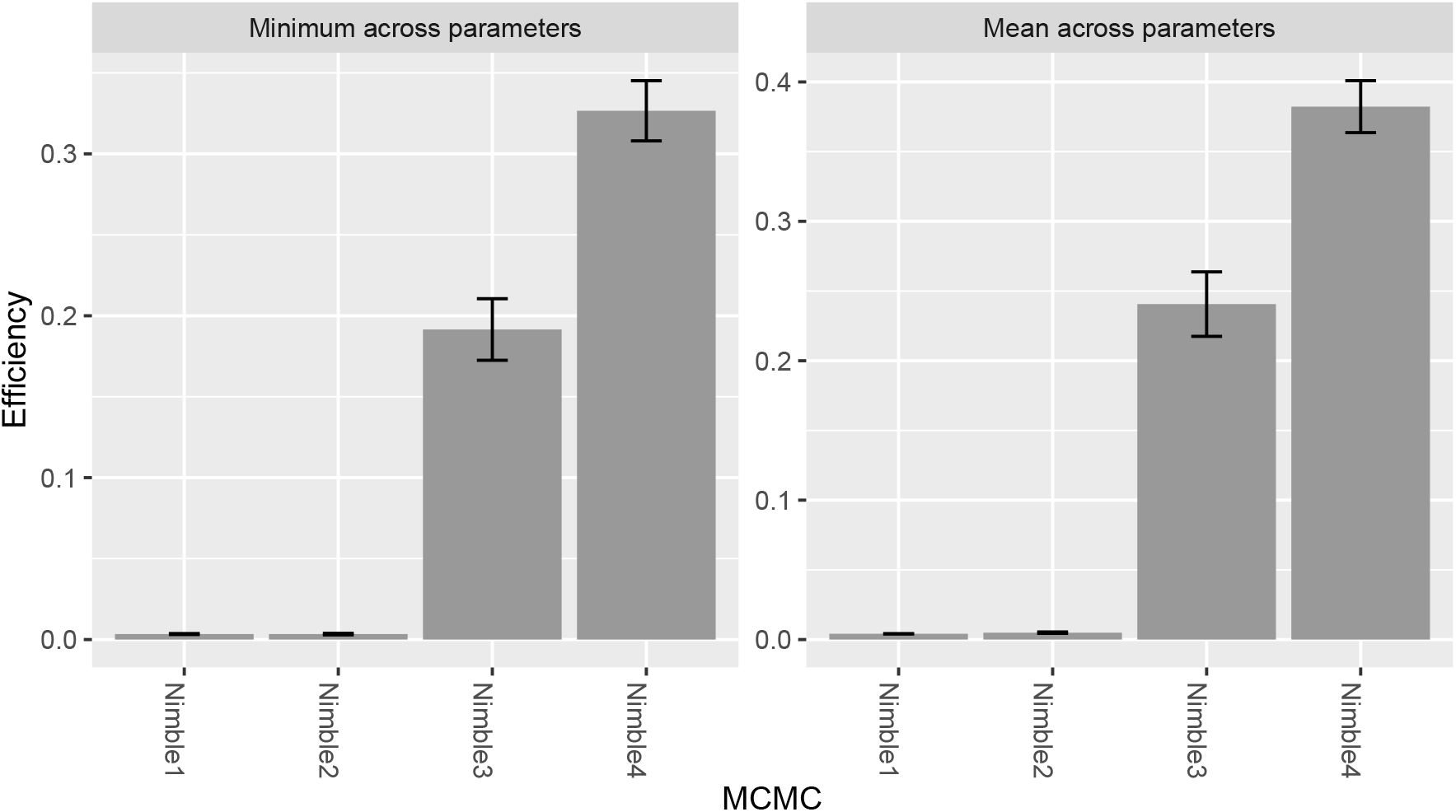
Minimum and mean MCMC efficiency among the three model parameters for the Wolverine example. Values are averaged over three independent chains, error bars showing standard deviation.

### 3.2 Vole Robust-Design Model

The Vole model contains a total of 11 hyper-parameters, which we use to assess MCMC efficiency. Results for the three stages of iterative improvement described in Section 2.2 will be denoted as Nimble1 (marginalization), Nimble2 (customizing dispersal distribution), and Nimble3 (evaluating local detectors). The original formulation of the model, running in JAGS, required over 4.5 hours to generate 10,000 posterior samples, and resulted in a minimum MCMC efficiency of 0.002, and a mean efficiency of 0.04.

The Nimble1 version introduced joint sampling of correlated parameters, and a custom likelihood to remove the *z*_*ij*_ latent states. This reduced the total model size from 4,460 nodes down to 3,562, while the number of unobserved nodes undergoing MCMC sampling was reduced from 1,437 down to 1,067. This model yielded an MCMC runtime of 15 minutes. ESS values were higher than those of JAGS, particularly for the jointly-sampled *σ*_*i*_ and *κ*_*i*_ parameters. MCMC efficiency was therefore higher for all parameters (Figure 3), while the average efficiency increased by a factor of 7.5 relative to JAGS.

**Figure 3:**
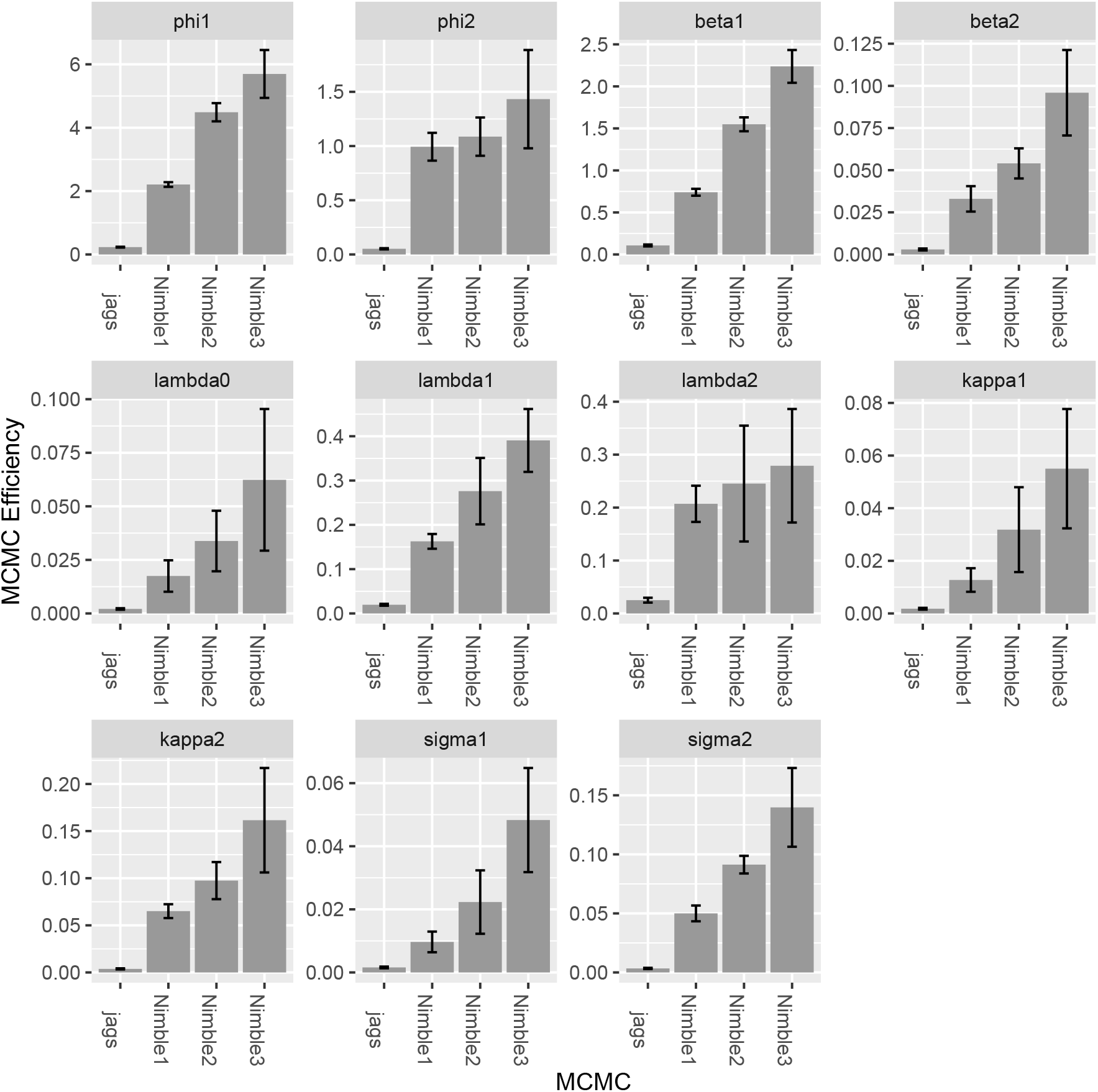
MCMC efficiency for the Vole example, for all parameters and model formulations. Efficiency values are averaged over five independent chains, error bars showing standard deviation.

The Nimble2 model introduced a custom bivariate dispersal distribution for individual ACs. This reduced the total model size from 3,562 nodes to 2,452, and the number of nodes for MCMC sampling from 1,067 to 697. Runtime decreased by a factor of two, to seven minutes, and all parameter MCMC efficiencies increased.

Using local trap calculations in the Nimble3 model reduced MCMC runtime further, to five minutes. Overall, relative to the initial analysis appearing in Ergon and Gardner (2014), these strategies reduced MCMC runtime by more than a factor of 50, and increased both minimum and mean MCMC efficiencies 25-fold. Concretely, the original model fitted in JAGS would require over seven days to produce 1,000 ESS for all parameters, whereas the Nimble3 formulation requires less than 6 hours to accomplish the same.

Figure 4 presents the minimum and mean efficiencies across all model parameters for each formulation of the Vole model, and all results for the Vole example appear in Table 4.

**Figure 4:**
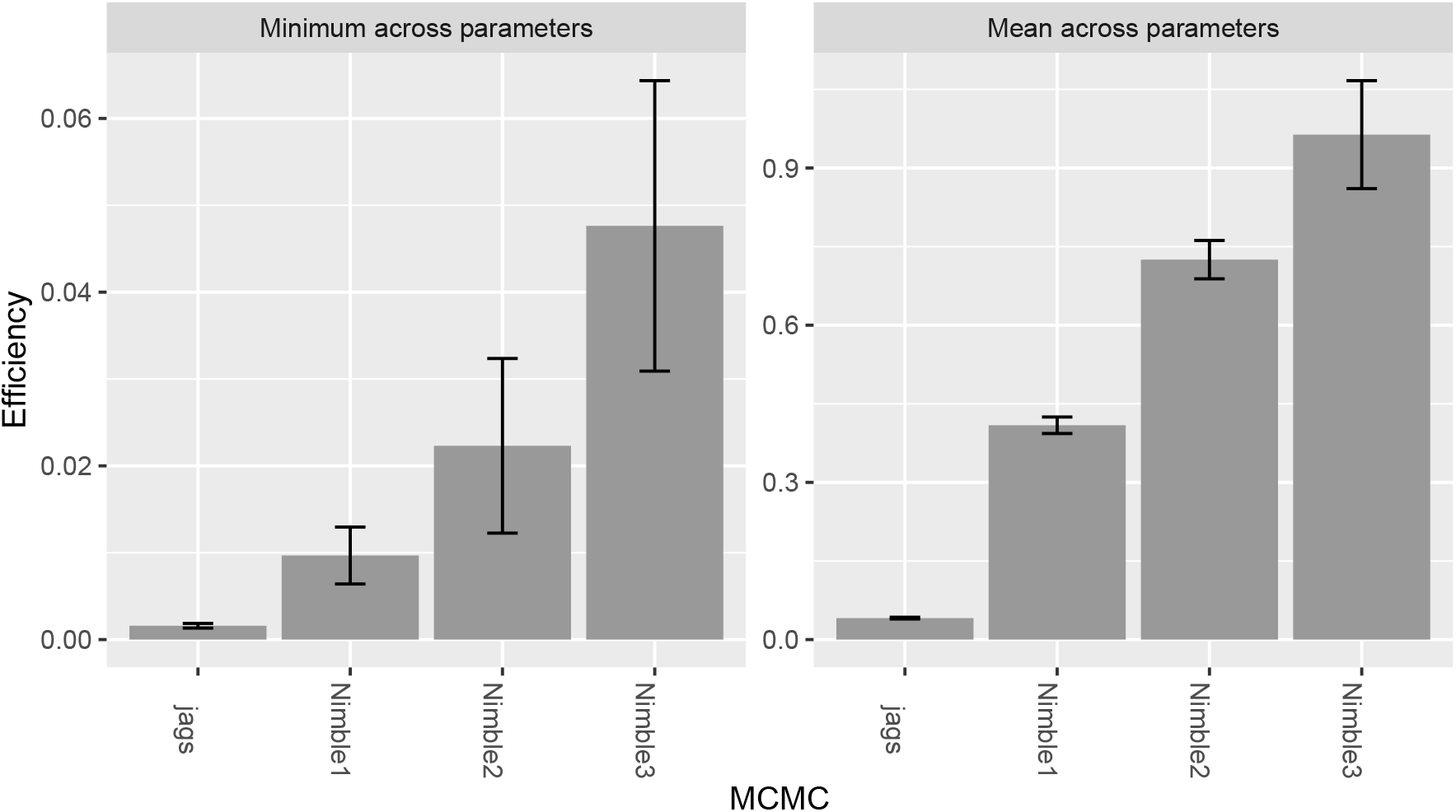
Minimum and mean MCMC efficiency among the eleven model parameters for the Vole example. Values are averaged over five independent chains, error bars showing standard deviation.

## 4 Discussion

SCR models are now commonplace given the abundance of geolocated ecological data, but remain computationally challenging. Indeed, large numbers of individuals, expansive study areas, and/or movement between seasons can render some problems intractable, without employing custom approaches.

The techniques demonstrated here produce posterior results identical (within Monte Carlo error) to the original versions, with the exception of local trap evaluations. This attributed a small trap hazard rate (or probability of detection) outside a radius *d*_min_ from each individual AC. The choice of *d*_min_ is important: large values will produce identical inferences but offer no computational gain, while small values offer a large computational gain but may introduce bias. The choice of *d*_min_ is subjective, and will require expert opinion (or trial runs) to determine an appropriate value. Smaller values of *d*_min_ may be used for exploratory analyses, but a conservative higher value should be used to minimize any biases in the final inference.

We are aware that conditioning on the primary occasion of first capture, as in the Vole example, may induce bias into parameter estimates (Efford and Schofield, 2019, Appendix E). Simulations in Ergon and Gardner (2014) suggest minimal bias in mortality estimates, although the scale parameter in the observation model may be inflated. Thus, care should be taken when applying this model to other data. That said, our purpose has been to investigate efficiency of estimation methods rather than statistical properties (such as bias or goodness of fit) of paticular models. Indeed, the ability to perform inference more efficiently will support a deeper exploration of alternative models structures.

Many software packages are available for fitting SCR models, making these analyses faster and more accessible to practitioners (*e.g.*, `secr`, or `oSCR`, among others). The prevalence of specialized software underscores the complex nature of SCR problems, and furthermore that no single software package could be general enough to approach all SCR problems. `nimble` does not attempt to provide “canned” algorithms for SCR, or any other particular application, but rather a flexible programming environment suitable for customized (and highly efficient) analysis of complex data.

We have made use of the `nimble` software package for `R`, to demonstrate techniques for improving the performance of SCR model fitting using MCMC. The techniques demonstrated are not exhaustive, but rather suggest the potential performance gains made possible using `nimble`, where we observed between one and two orders of magnitude improvement. These approaches can provide significant computational gain, permitting large-scale spatial and temporal analyses to support major conservation and management decisions, and the ability to fit increasingly complex models to large datasets. More broadly, similar techniques are also applicable to the analysis of general spatially-indexed hierarchical model structures.

**Table 1:** MCMC efficiency values for the Wolverine example, for all parameters and model formulations. Results are averaged over three independent chains.

|  | Parameter |  |  |
| --- | --- | --- | --- |
| | $N$ | $p_0$ | $\sigma$ |
| Nimble1 | 0.005 | 0.004 | 0.003 |
| Nimble2 | 0.006 | 0.005 | 0.003 |
| Nimble3 | 0.284 | 0.247 | 0.192 |
| Nimble4 | 0.390 | 0.394 | 0.362 |

**Table 2:** MCMC efficiency values for the Vole example, for all parameters and model formulations. Results are averaged over five independent chains.

|  | Parameter |  |  |  |  |  |  |  |  |  |  |
| --- | --- | --- | --- | --- | --- | --- | --- | --- | --- | --- | --- |
| | $\phi_1$ | $\phi_2$ | $\beta_1$ | $\beta_2$ | $\lambda_0$ | $\lambda_1$ | $\lambda_2$ | $\kappa_1$ | $\kappa_2$ | $\sigma_1$ | $\sigma_2$ |
| JAGS | 0.23 | 0.05 | 0.11 | 0.003 | 0.002 | 0.02 | 0.03 | 0.002 | 0.004 | 0.002 | 0.003 |
| Nimble1 | 2.21 | 0.99 | 0.74 | 0.033 | 0.017 | 0.16 | 0.21 | 0.013 | 0.065 | 0.010 | 0.050 |
| Nimble2 | 4.49 | 1.09 | 1.55 | 0.054 | 0.034 | 0.28 | 0.25 | 0.032 | 0.097 | 0.022 | 0.091 |
| Nimble3 | 5.70 | 1.43 | 2.24 | 0.096 | 0.062 | 0.39 | 0.28 | 0.055 | 0.162 | 0.048 | 0.140 |

## Acknowledgements

We would like to thank Pierre Dupont and Richard Bischof for their help with the Wolverine example analysis. This work was partly funded by the Norwegian Environment Agency (Miljødirektoratet), the Swedish Environmental Protection Agency (Naturvårdsverket) and the Research Council of Norway (NFR 286886).

## A Wolverine Example: Model Code

## A.1 Vectorized Computations Code

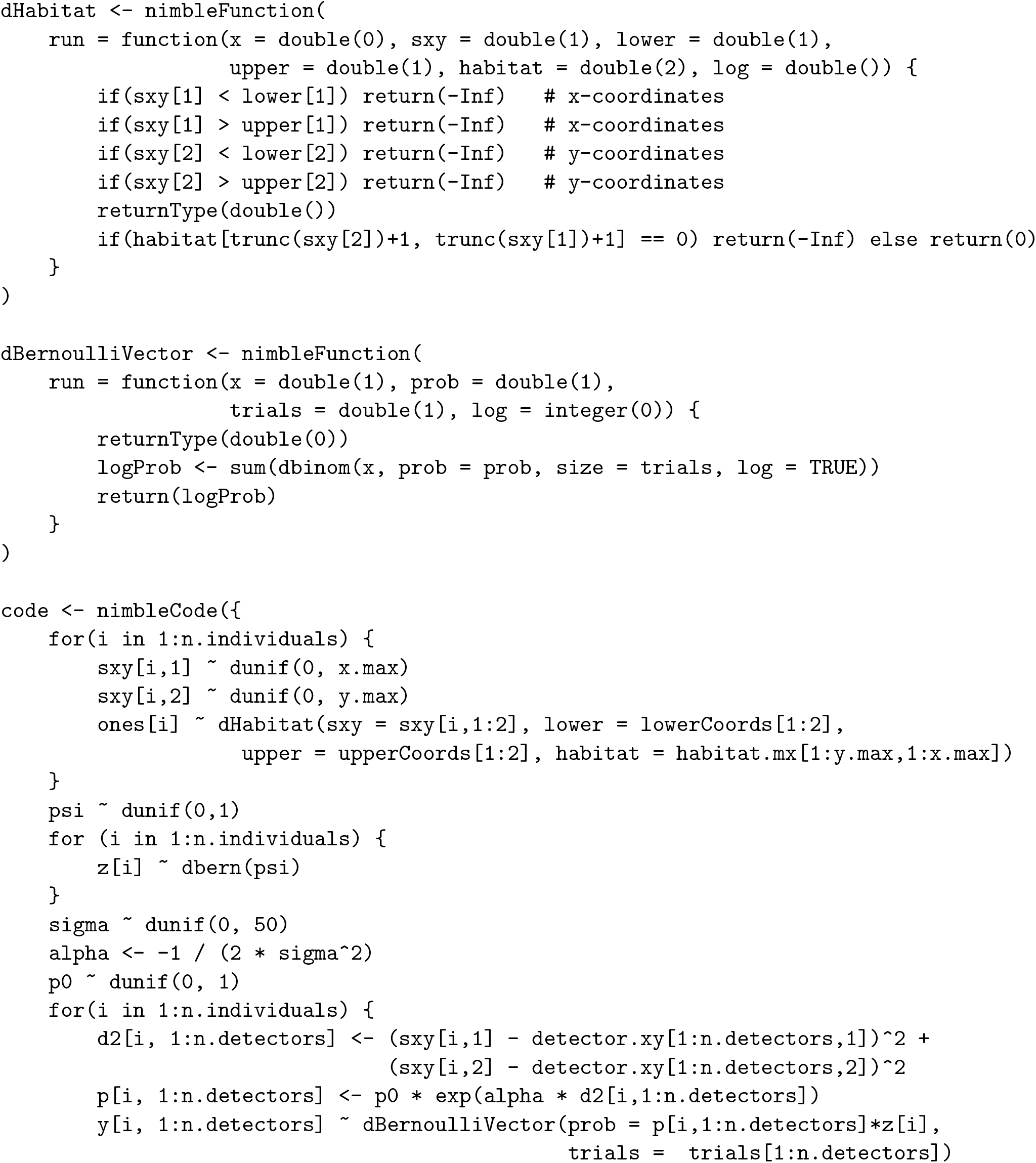
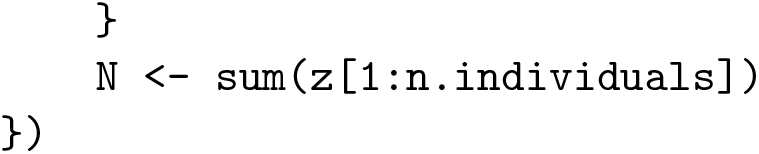

## A.2 Local Detector Evaluations and Sparse Observation Matrix Code

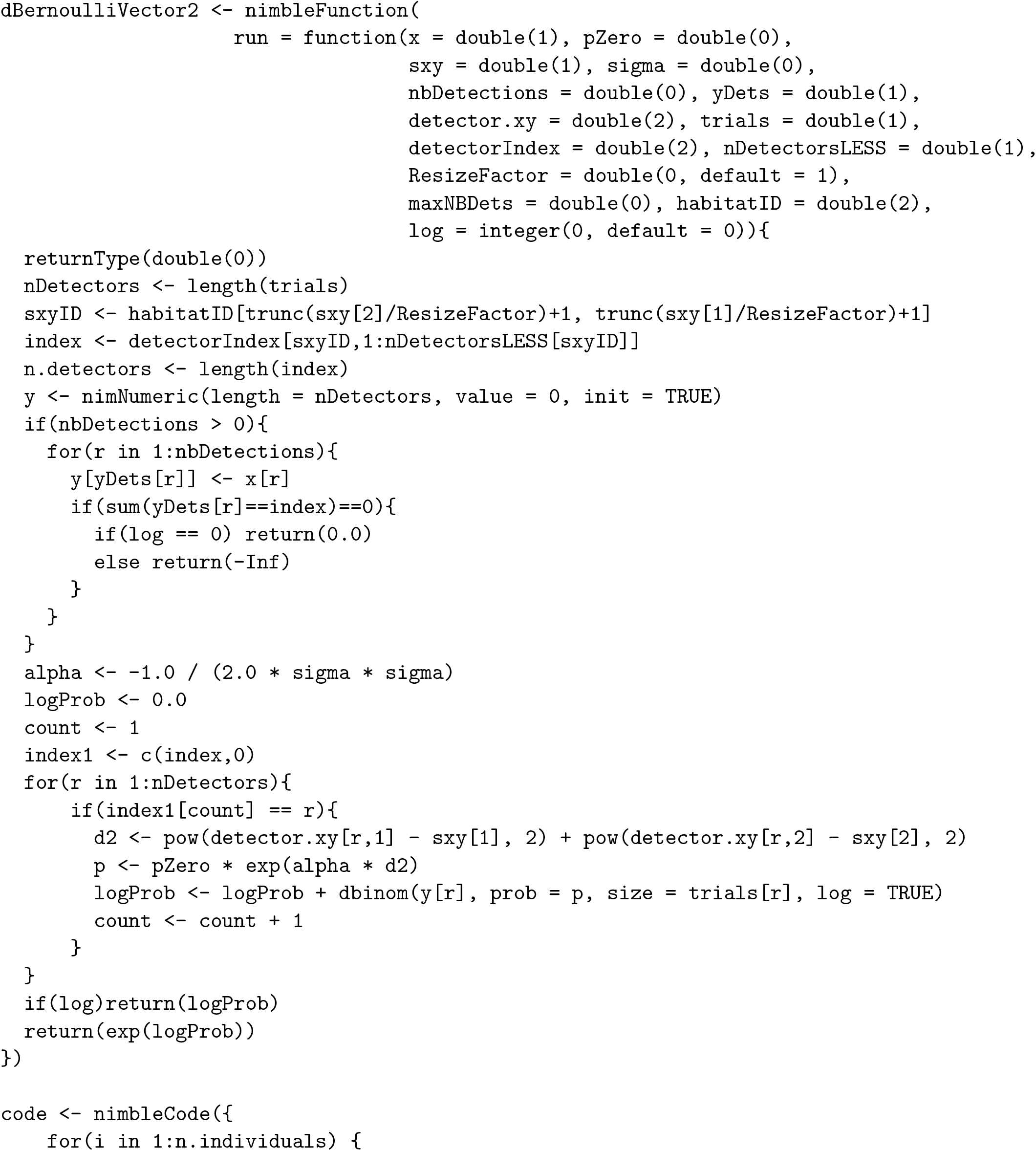
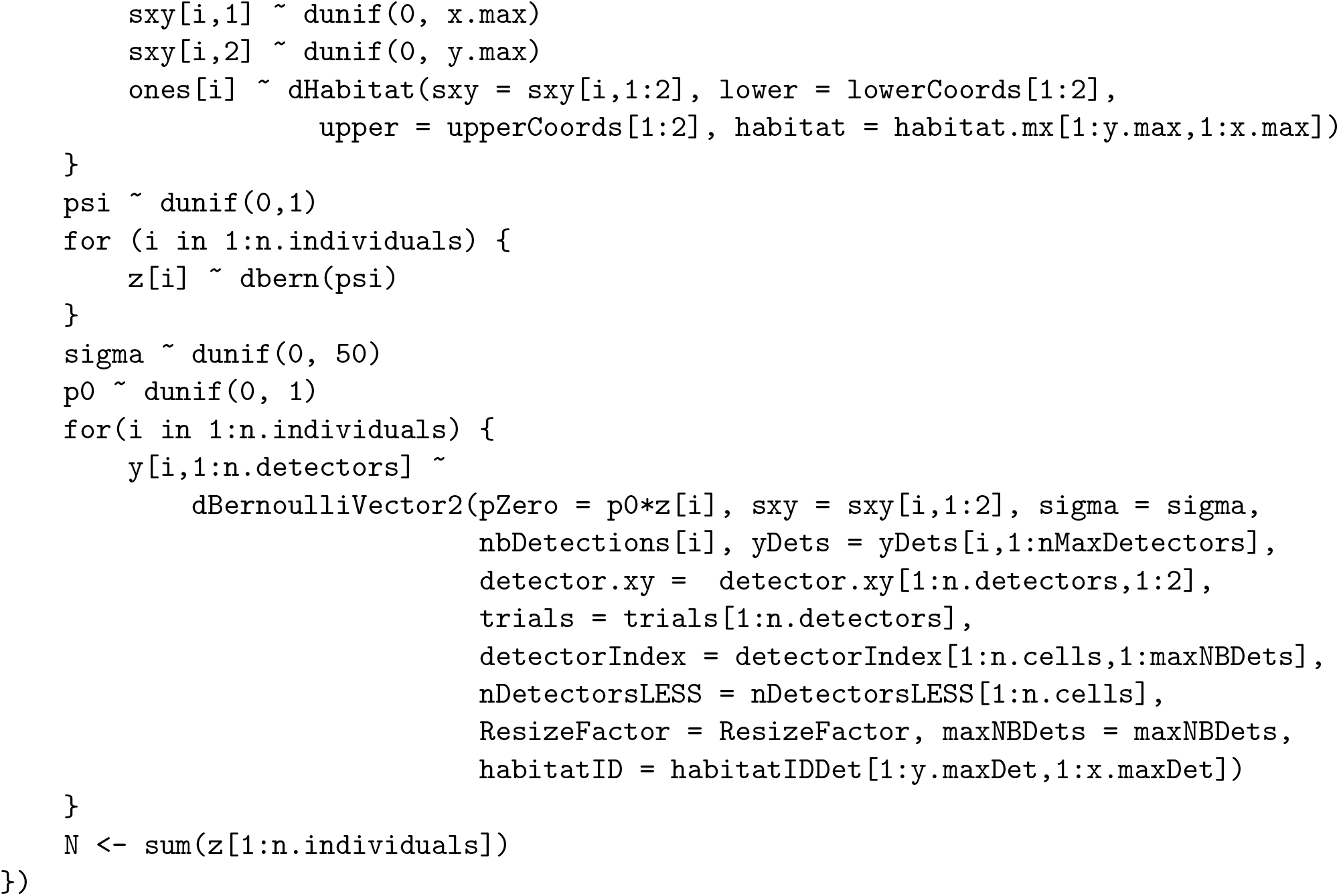

## A.3 Skip Local Calculations Code

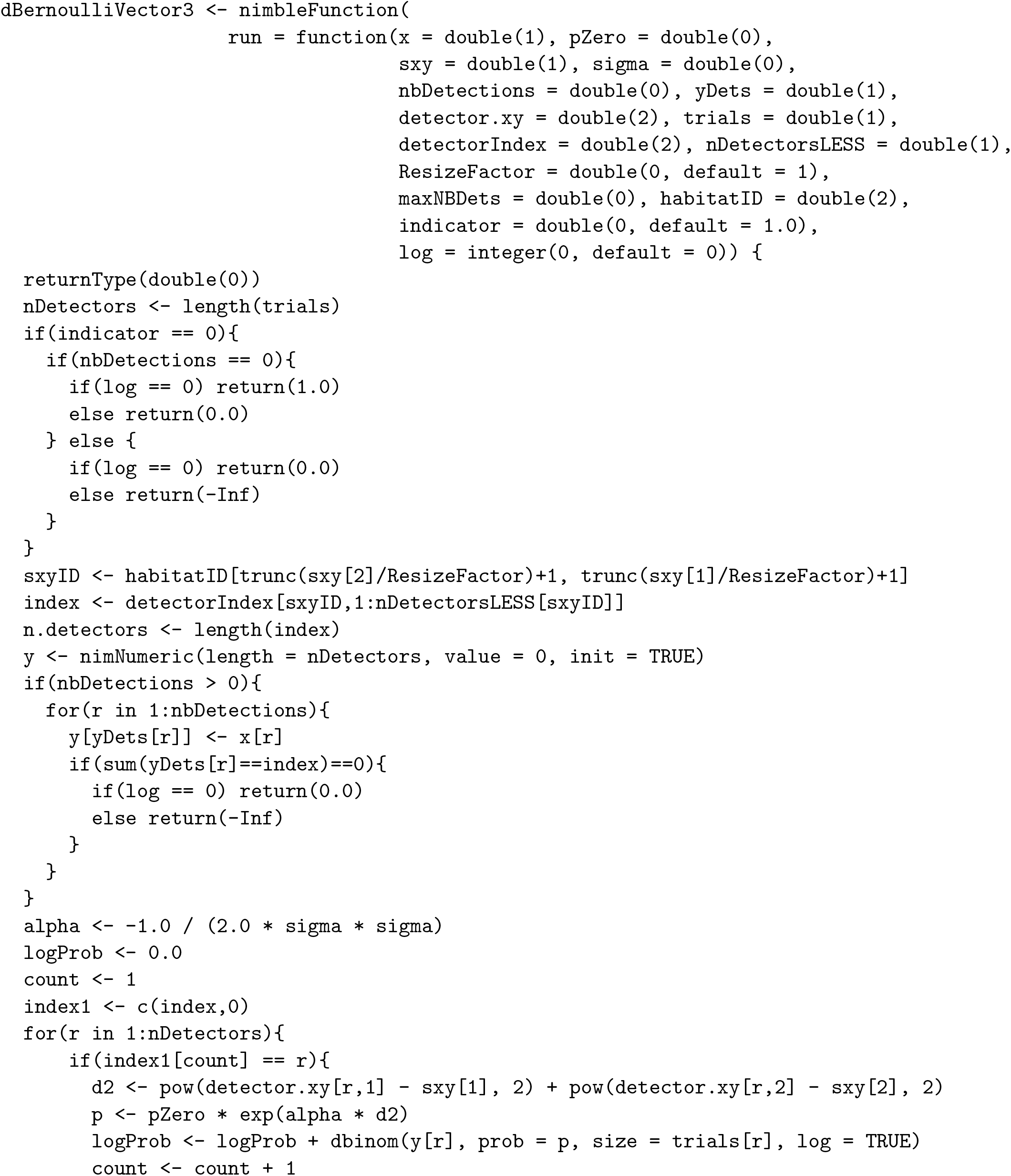
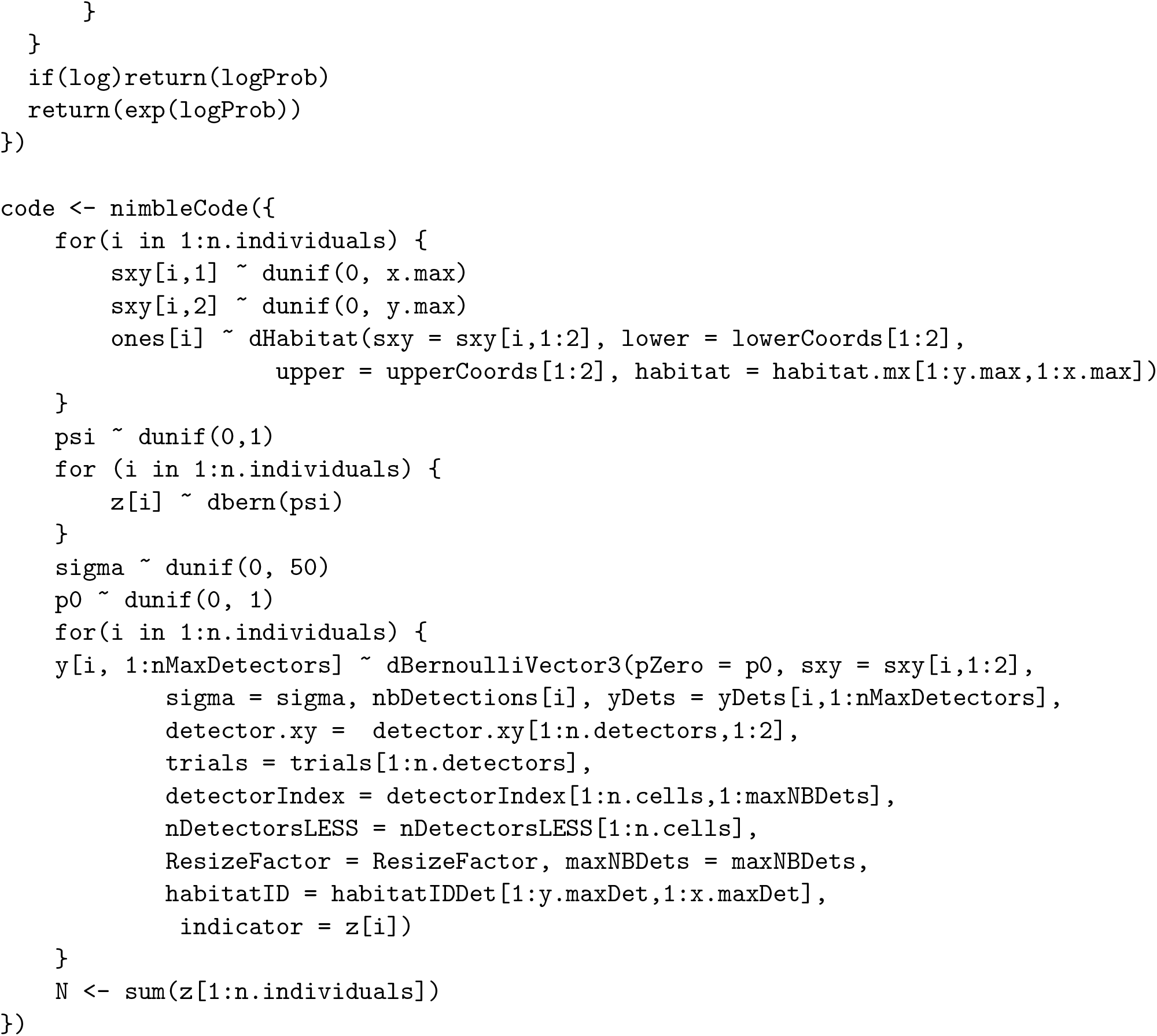

## B Vole Example: Model Code

## B.1 JAGS Code

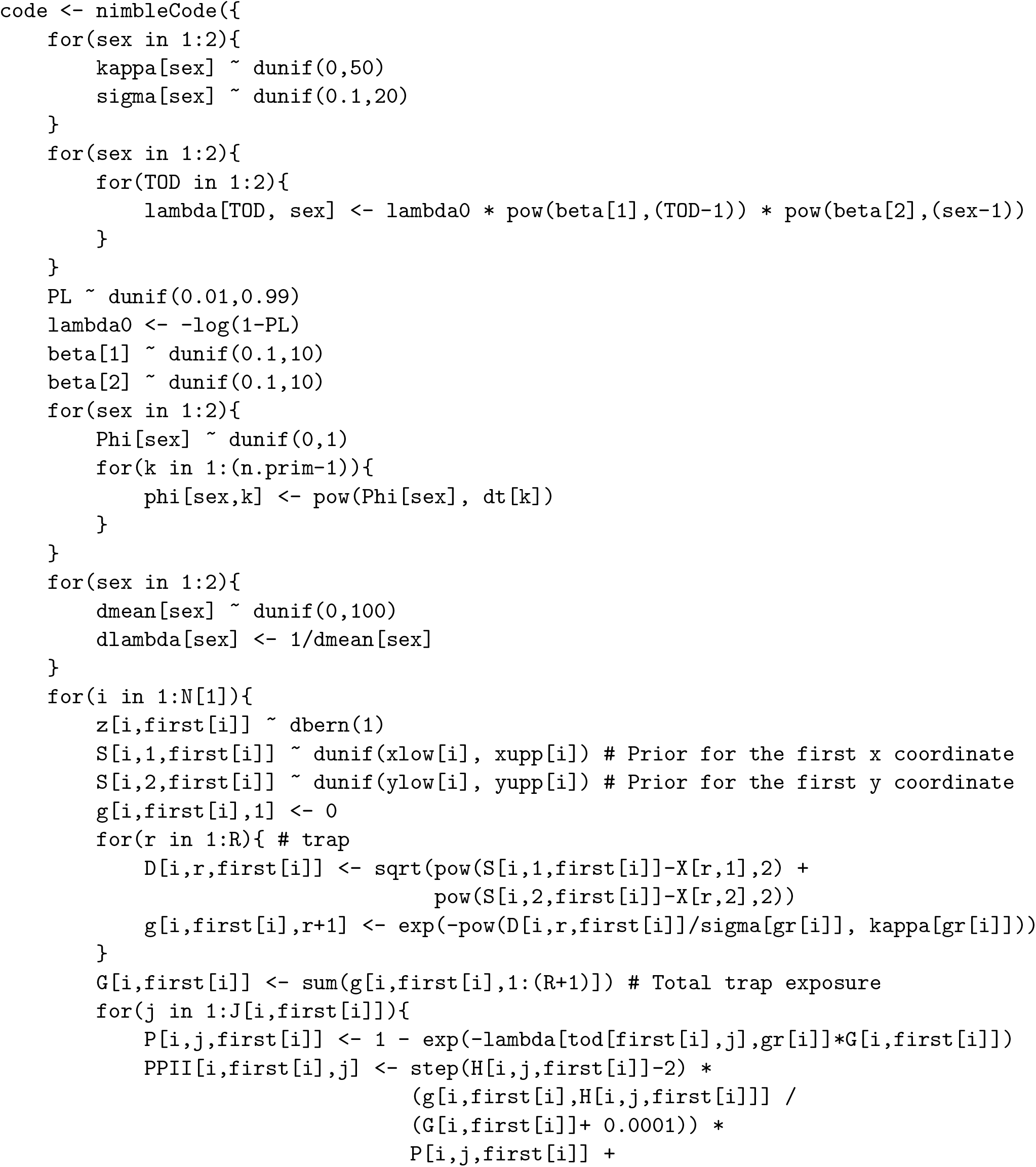
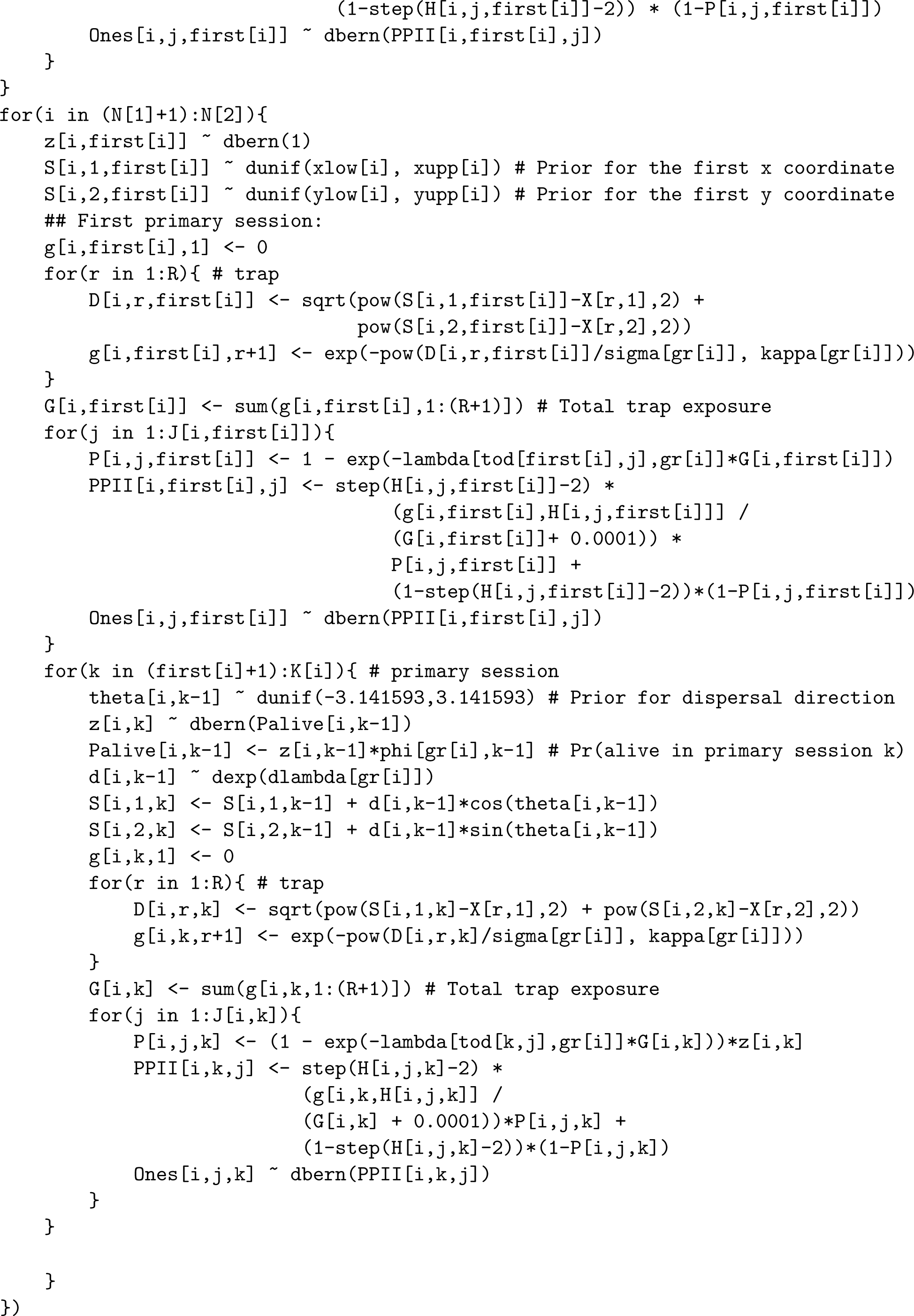

## B.2 Joint Sampling and Marginalization Code

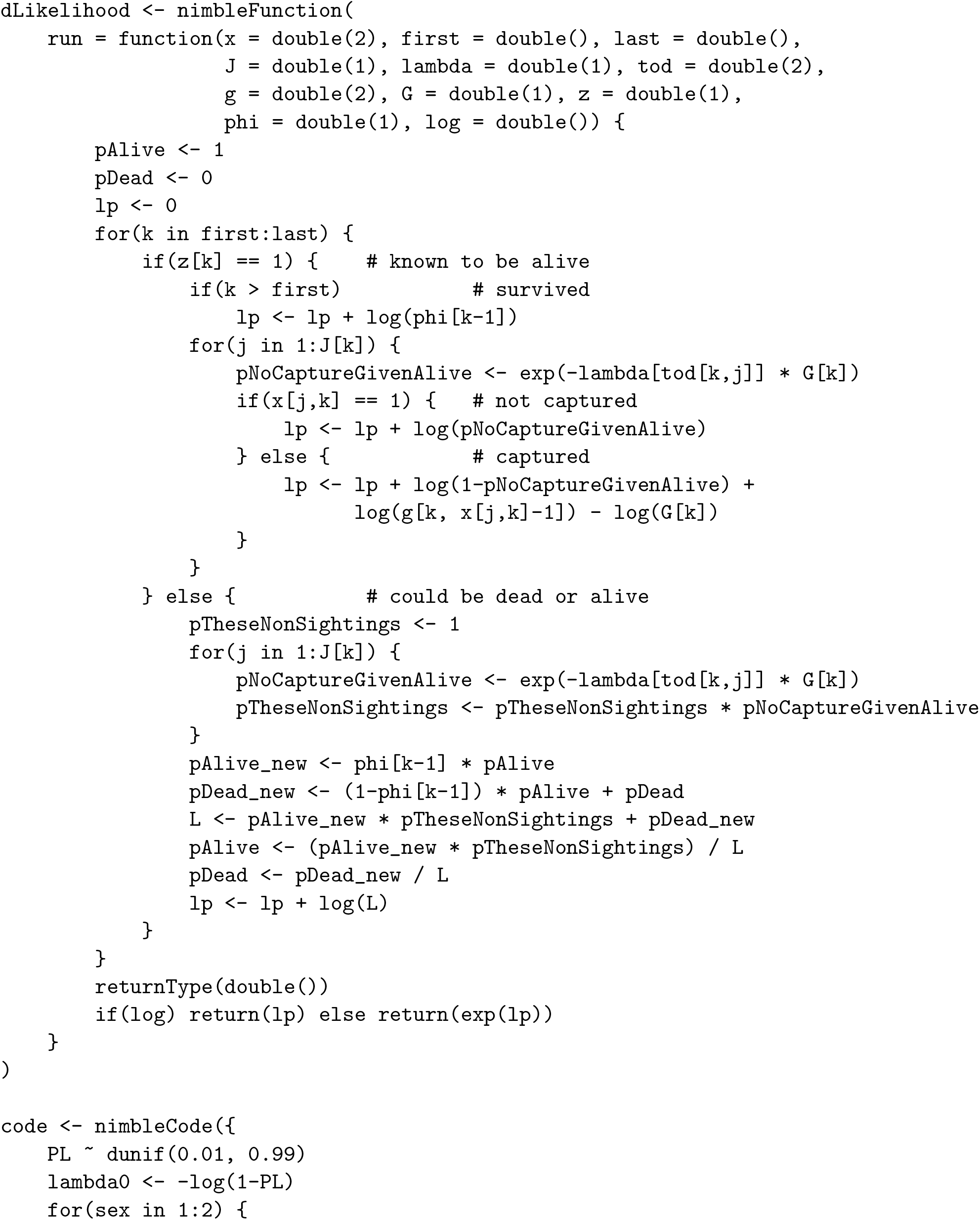
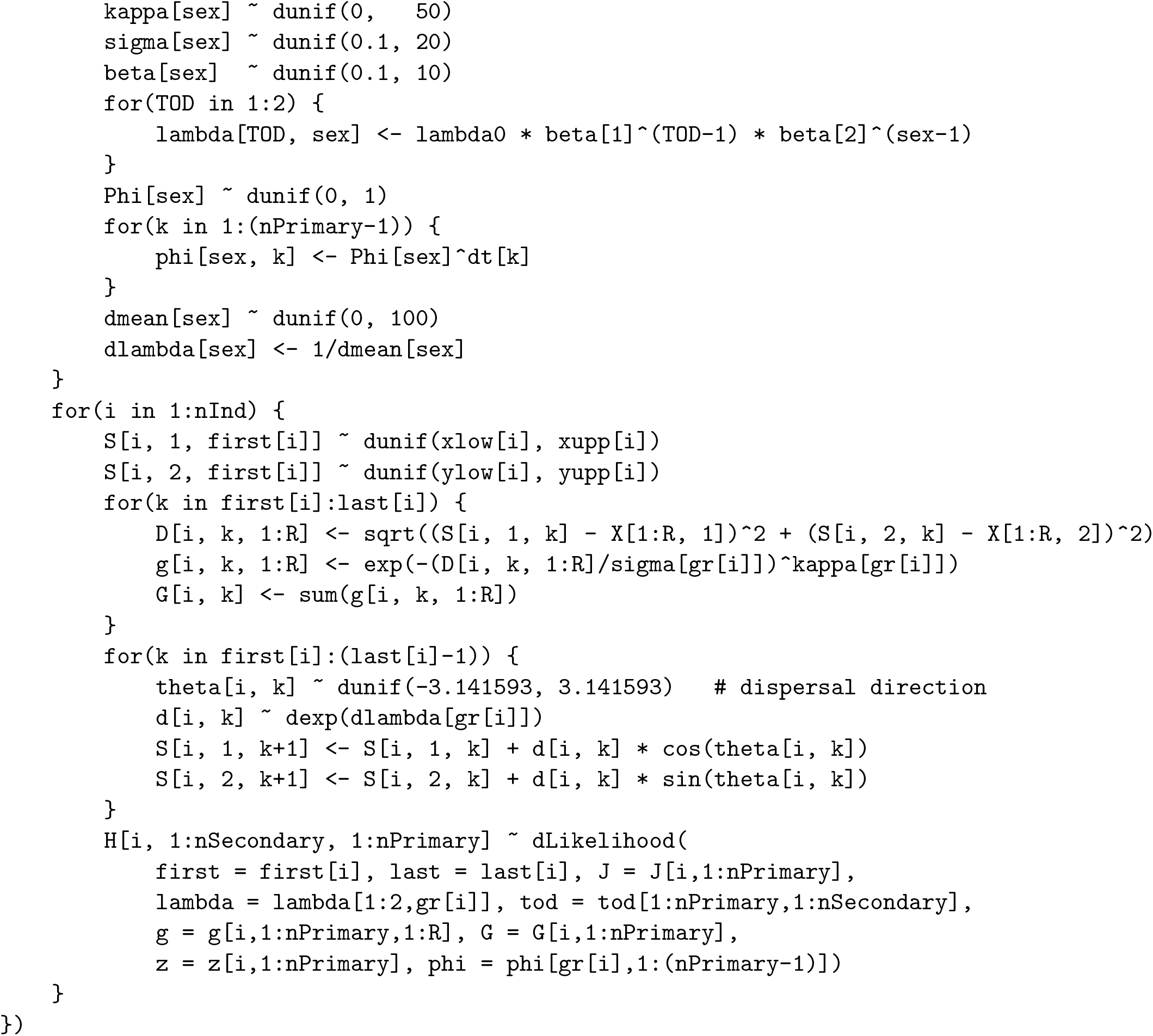

## B.3 Custom Dispersal Distribution Code

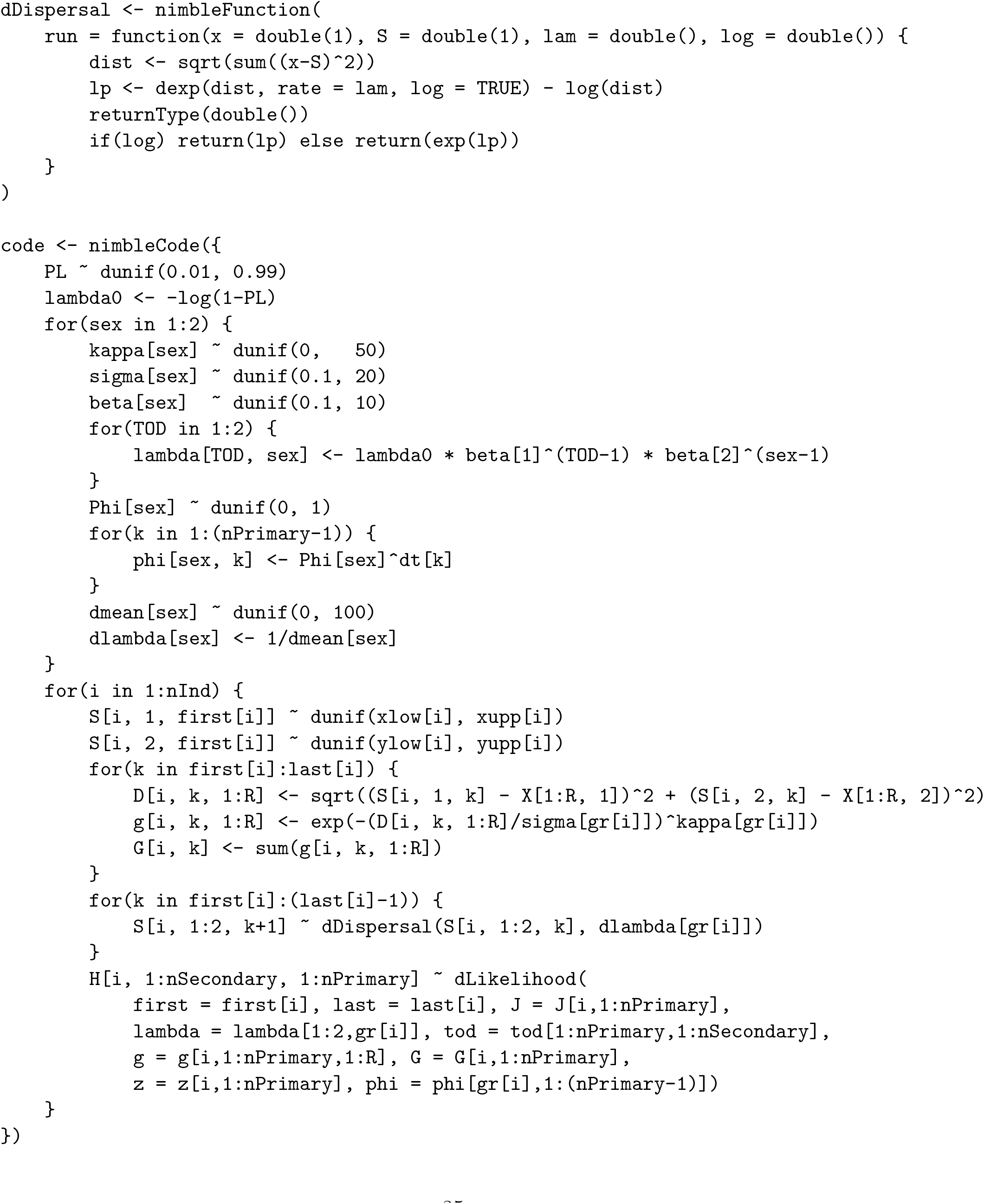

## B.4 Local Trap Calculations Code

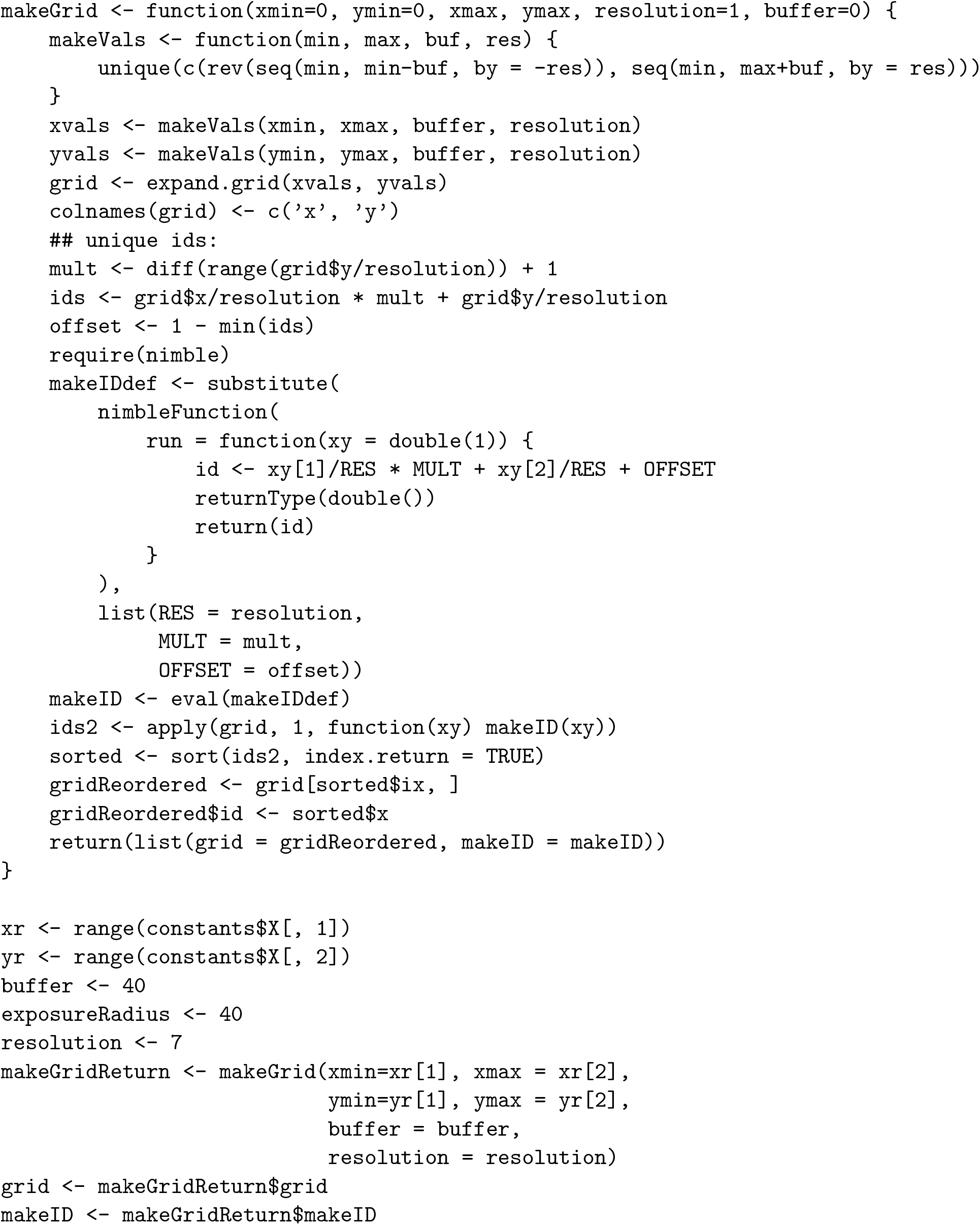
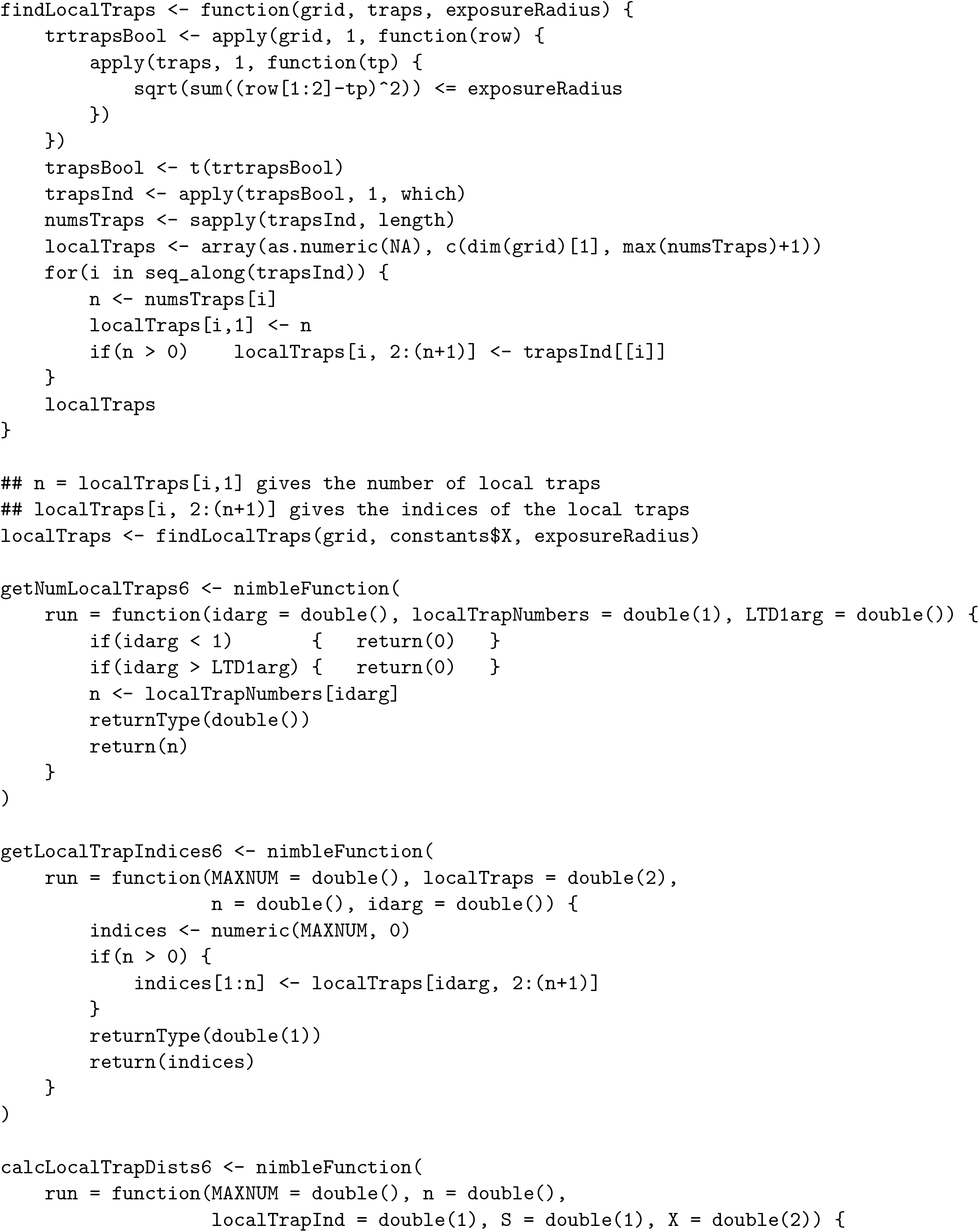
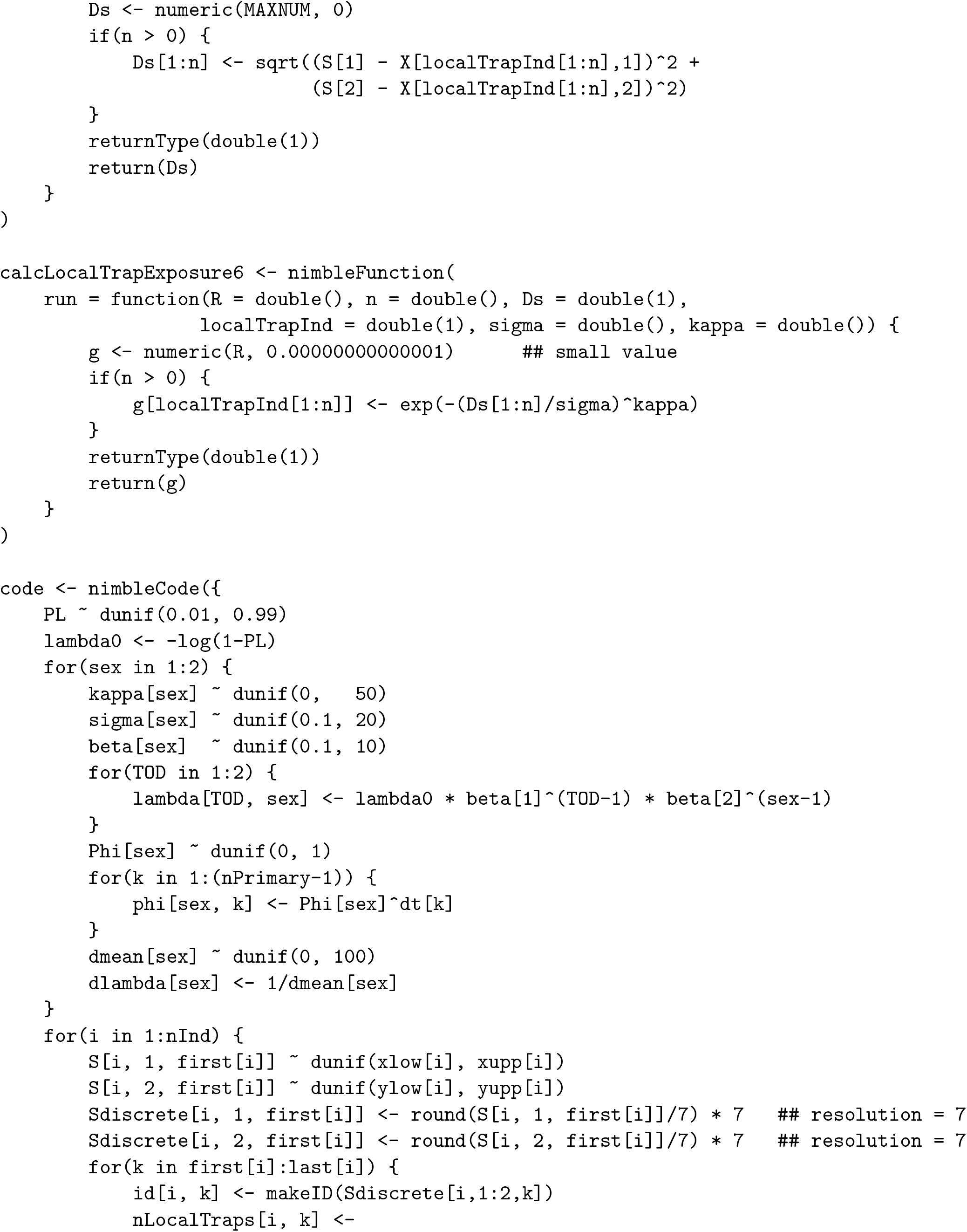
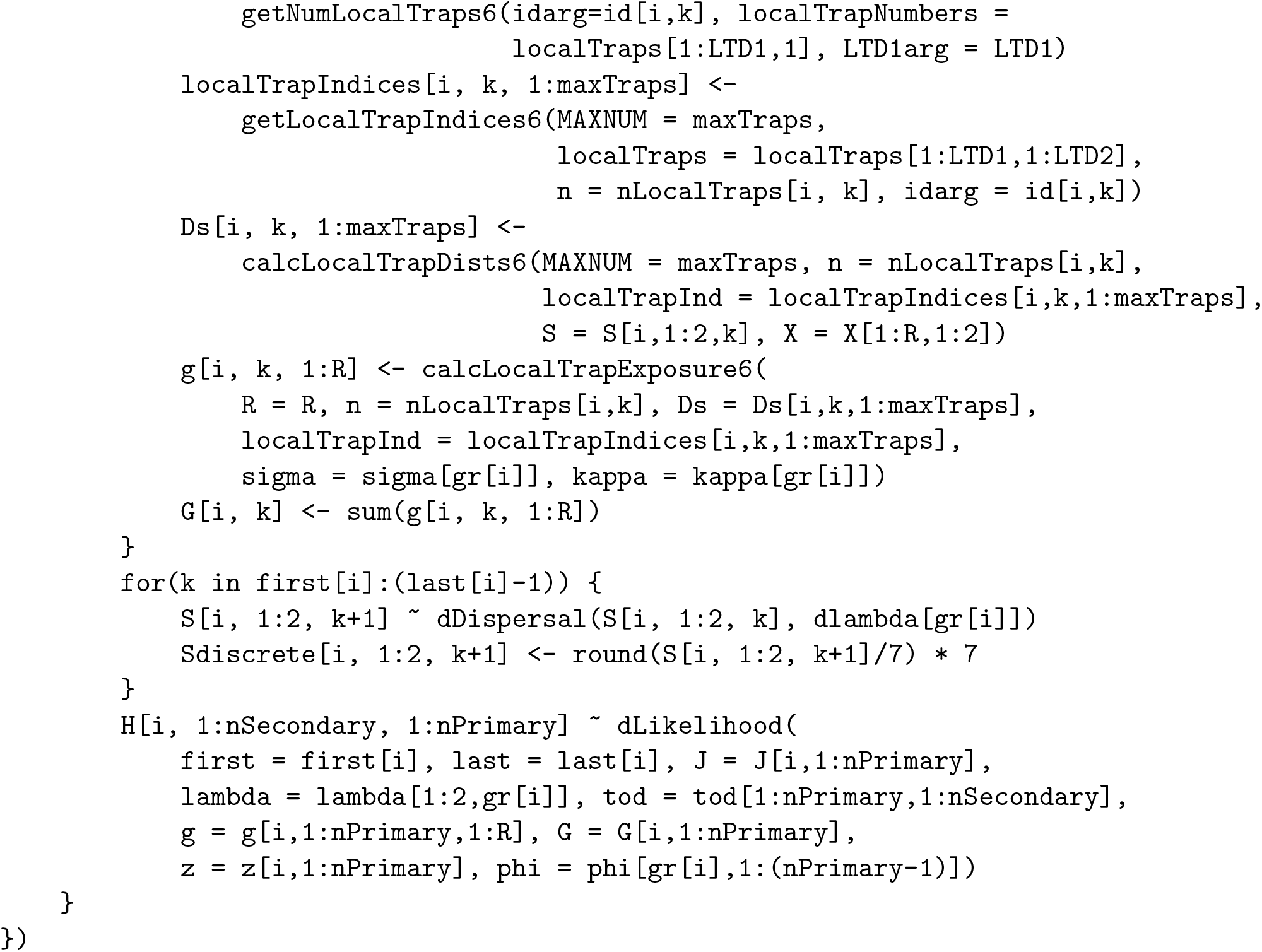

